# Imaging the Electrical Activity Of Organelles in living cells

**DOI:** 10.1101/578765

**Authors:** Ella Matamala, Cristian Castillo, Juan Pablo Vivar, Sebastian Brauchi

## Abstract

Eukaryotic cells are complex systems compartmentalized in membrane-bound organelles. Visualization of organellar electrical activity in living cells requires both a suitable reporter and non-invasive imaging at high spatiotemporal resolution. Here we present hVoS_org_, an optical method to monitor changes in the membrane potential of subcellular membranes. This method takes advantage of a FRET pair consisting of a membrane-bound voltage-insensitive fluorescent donor and a colorless voltage-dependent acceptor that rapidly moves across the membrane in response to changes in polarity. Compared to the currently available techniques, hVoS_org_ has advantages including simple and precise subcellular targeting, the ability to record from individual organelles, and the potential for optical multiplexing.

**Summary:** By adapting a hybrid-FRET voltage sensor we report here the resting membrane potential of different organelles in living cells.

## Main Text

In general, charge separation as a result of the regulated flow of ions establishes a voltage gradient across semipermeable membranes. The voltage gradient across organelle membranes (Ψ_org_) will be then defined by the specific set of ion channels and transporter proteins expressed on a given organelle. Ψ_org_ is modulated by intracellular signaling cascades and is likely to be essential to the maintenance of organellar homeostasis ^1–6^. While the central importance of the electrical activity of organelles has been widely acknowledged, the detailed mechanisms that support this type of signaling are poorly understood, partially due to the lack of the sophisticated research tools that have been developed for studies of voltage gradient changes at the plasma membrane ^7–11^.

Membrane potential imaging using voltage-sensitive dyes has been extensively used for mitochondria and ER; and more recently, for phagosomes and lysosomes ^12–15^. Still, a precise and standardized method allowing for the recording of electrical signals generated at individual organelles is not yet available. To this end, we have developed a general methodology for the recording of electrical signals from individual organelles. The method relies on the use of a Hybrid Voltage Sensor (hVoS) ^16^ and here we show its effectiveness for the fast imaging of variations in Ψ_org_ (∆Ψ_org_) in intact living cells.

The hVoS approach is extremely sensitive, capable of measuring rapid changes in the membrane potential of both excitable and non-excitable cells ^16–19^. The method takes advantage of a FRET pair consisting of a membrane-anchored fluorescent protein acting as donor and the colorless hydrophobic anion dipicrylamine (DPA) acting as acceptor. Due to its small size, negatively charged DPA has the ability to rapidly transit across the membrane in response to changes in the membrane potential, acting as voltage sensor ^16^. Conveniently, the imaging read-out of membrane potentials at the plasma membrane is linear within a broad dynamic voltage range (from −130 to +40 mV) ^18^.

In principle, when combined with DPA, it is possible for any membrane-bound fluorescent marker to transduce voltage changes occurring at the target membrane into fluorescence fluctuations. Therefore, fluorescently-tagged protein markers that are routinely used for selective subcellular expression provide a handy tool to image the electrical activity of internal membranes. For hVoS_org_ to work, DPA must reach intracellular membranes (Fig. 1a). Once at the membrane it would distribute according a voltage-dependent equilibrium ^18,20^. In such a scenario, each individual membrane compartment of the cell would define three equilibriums governing the distribution of DPA molecules – a lipid-water interphase at each side of the membrane and a voltage-dependent barrier governing DPA transit between the them (Fig. 1a) ^21^.

**Figure 1.**
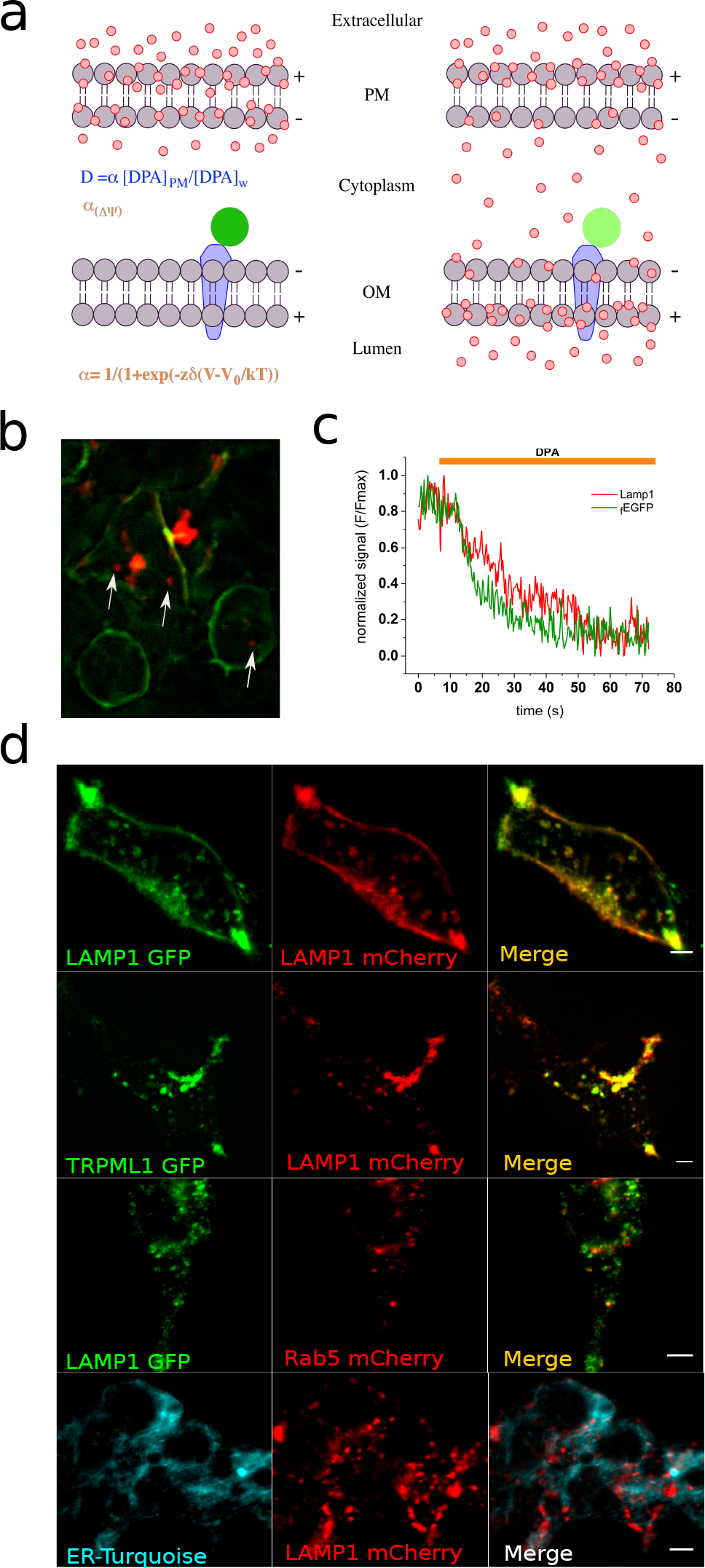
Dipikrylamine (DPA) reaches internal membranes. **(a)** Schematic representation of DPA incorporation and distribution within the cell. Knowing the concentration of DPA at the extracellular medium and the partition coefficient (*D*), it is possible to calculate the concentration at the membrane and also the voltage-dependent probability (*α*) of being on a given leaflet. The two membranes are in series and are opposite in polarity, causing accumulation of DPA in the lumen of internal compartments. The voltage-dependent movement of DPA molecules within the membrane will alter GFP fluorescence, making us able to differentiate hyperpolarization and depolarization of the organelle. **(b)** Simultaneous expression of Lamp1-mCherry (*lysososme marker*) and farsenylated GFP (*plasma membrane marker*) in HEK293 cells. Arrows indicate lysosomes not forming clusters. **(c)** Time course of fluorescence for the cell in b when exposed to DPA. The lysosomal signal corresponds to the average of the three spots indicated by arrows in panel b. **(d)** Representative colocalization images of the lysosomal marker Lamp1. Similar level of colocalization are observed for EGFP-/mCherry-tagged Lamp1 and Lamp1-mCherry / TRPML1-GFP. A low level of colocalization is observed between the endosomal marker Rab5 and Lamp1 or between the ER marker ER-turquoise/Lamp1-EGFP. Scale bars correspond to 5 µm.

Lysosomes are degradative organelles essential to maintain cellular metabolic activity. Their direct association with mTOR kinases is thought to integrate their catabolic role with different signaling cascades in the cell ^22^. Several channels, transporters, and ion pumps such as the vesicular proton pump (v-ATPase), Two-Pore Na^+^ Channels (TPCs), TMEM175 K^+^ channels, members of the mucolipin subfamily of TRP channels (TRPMLs), SLC, CLC, and CLIC transporters, have been described as active residents of the endolysosomal system ^22–24^. The expression patterns and localization of these proteins combined with electrophysiological data has led to the proposal that the lysosome is an electrically active organelle ^6,15^.

Imaging of the membrane potential of lysosomes (Ψ_ly_) have been previously accomplished by using a combination of potentiometric fluorescent dyes (i.e. oxonol derivatives) forming a FRET pair with fluorophores that are *preferentially* segregated to the lysosome membrane ^15^. However, the high density and variety of membrane-bound organelles within the endolysosomal system makes it impossible to isolate the individual contribution of endosomes, lysosomes, or phagosomes when a FRET pair is formed by hydrophobic dyes, imposing restrictions on spatial resolution and targeting.

Therefore, to optically follow rapid changes in Ψ_ly_, we recorded voltage-sensitive FRET signals between DPA and a fluorescent protein of choice (i.e. EGFP or mCherry) fused to the cytoplasmatic C-terminal domain of the lysosomal-associated membrane protein 1 (Lamp1). Our results indicated that hVoS_org_ reliably reports the amplitude and kinetics of Ψ_ly_ at the level of single organelle. Being a single wavelength excitation/emission tool, based on cellular markers of common use, the versatility of the technique allowed us for out-of-the-box recordings of other intracellular compartments. We report the resting potential of Golgi and ER membranes as examples of whether the technique could be easily expanded and multiplexed.

## Characterization and targeting of hVoS_org_ to lysosomal membranes

To first confirm that DPA can reach organellar membranes, we simultaneously followed two fluorescent markers, a farsenylated EGFP (_f_EGFP) that targets to the plasma membrane and Lamp1-mCherry that targets to the lysosome (Fig. 1b). As expected, we observe a loss of both fluorescent signals upon DPA addition (4 µM) and that the quenching of the signal at the plasma membrane precedes the response at the lysosome (Fig. 1c). This confirms the effectiveness of the intracellular FRET pair, and more importantly, the ability of DPA to reach the internal membranes in living cells. According to previous studies, the FRET efficiency of hVoS is larger for fluorescent proteins that are excited at lower wavelengths ^18^. Accordingly, most optical measurements of ΔΨ_ly_ were performed using Lamp1-EGFP to benefit the signal-to-noise ratio of the readout.

Lamp1 fluorescent constructs display a characteristic distribution that space-correlates well with other lysosome-resident proteins such as TRPML1 (mucolipin) and two-pore sodium channel 1 (TPC1) showing Pearson’s coefficients of 0.902 ± 0.047 and 0.894 ± 0.063 respectively (Fig. 1d). In contrast, the endosomal marker Rab5 and the ER marker mTurquoise2-ER do not show significant colocalization with Lamp1 (Fig. 1d).

In order to interpret the FRET readout, it is critical to know the orientation of the membrane-anchored fluorescent protein within the organelle membrane. To address this, we performed a fluorescence protease protection (FPP) assay ^25^. According to the protein’s design, the EGFP domain of our lysosomal marker should face to the cytoplasm. In good agreement to this, EGFP fluorescence rapidly quenches with trypsin (4 mM) after gentle permeabilization of the plasma membrane with digitonin (10 µM) (Fig. 2a, *top*). We further validated this result with a fluorescence quenching experiment using Lyso-pHoenix, a lysosome-targeted construct expressing the pH sensor pHluorin at the luminal side and the red-emitting fluorescent protein mKate facing the cytoplasm ^26^. Under the same conditions used for Lamp1-EGFP, pHluorin signal remains stable while mKate fluorescence rapidly quenches upon the addition trypsin (Fig. 2a, *bottom*).

**Figure 2.**
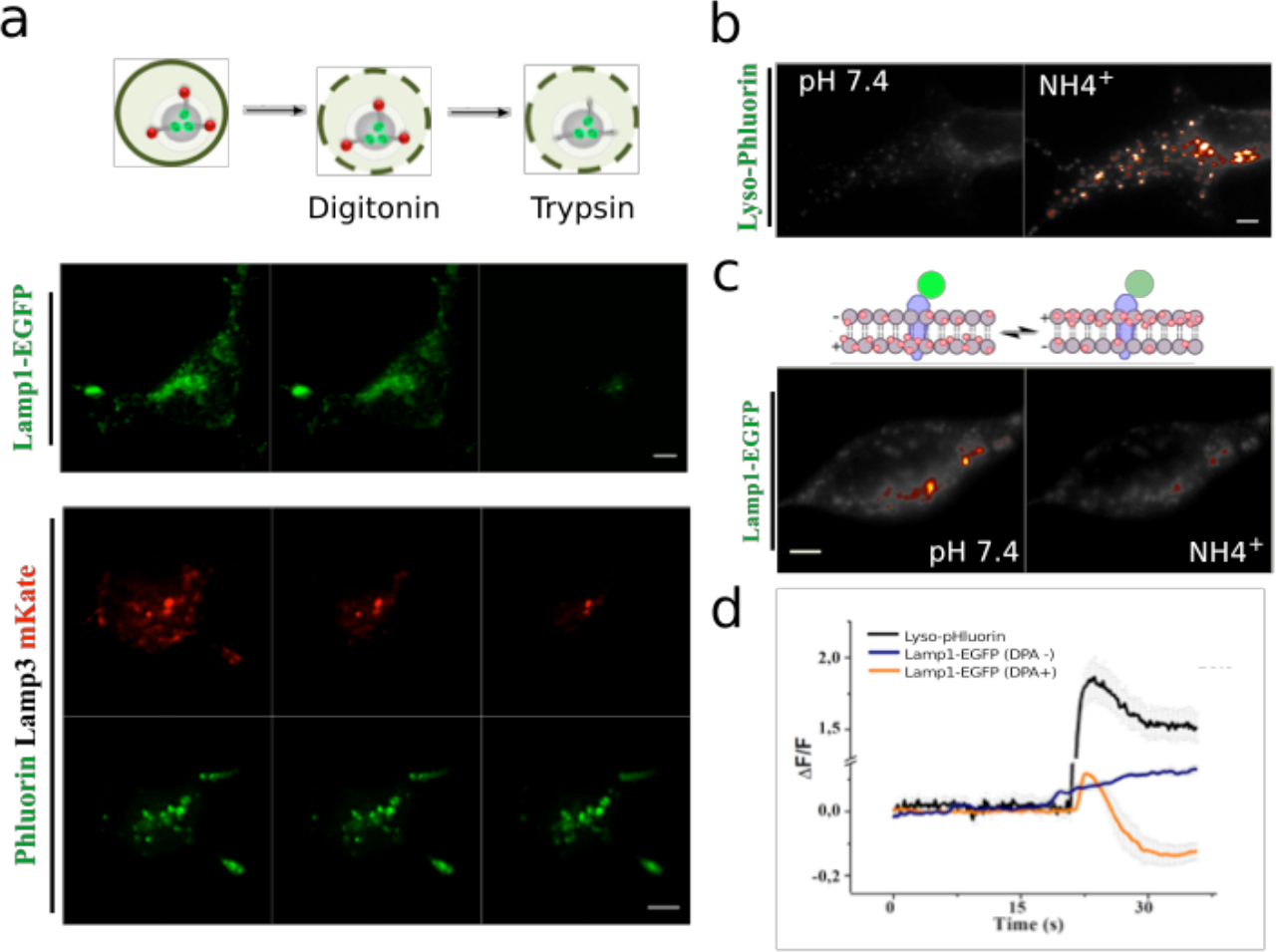
Topology of Lamp1-EGFP by FPP assay and the effect of lysosomal pH on hVoS_org_ signals. **(a)** Cartoon of the FPP assay illustrating the position of the fluorescent tags relative the organelle membrane (*top*). Lamp1-EGFP exposes the fluorescent protein to the cytoplasm as revealed by protease protection assay (scale bar, 5 µm). HEK-293T cells expressing lyso-pHoenix (pHluorin-CD63-pHluorin-Arch3-mKate) were subjected to the FPP assay (*lower panels*). FPP in this case revealed that pHluorin faces the lumen while mKate faces the cytoplasm. Images were taken before and after 2-min treatment with 10 µM digitonin followed by perfusion with a ringer containing 4 mM trypsin. The steps of the assay are indicated in the cartoon on top. **(b)** Fluorescence images of HEK-293 cells transfected with a plasmid encoding for a luminal pH sensor (pHluorin), before (*left*) and after (*right*) ammonium treatment. **(c)** Fluorescence images of HEK-293 cells transfected with a Lamp1-EGFP), before (*left*) and after (*right*) ammonium treatment. The cartoon on top depicts the transit of DPA molecules (red circles) and quenching of EGFP’s fluorescence (green circles) during depolarization of the lysosome. **(d)** Plot if intensity versus time for the ammonium treatment in different conditions. The arrow denotes addition of NH4^+^.

Wide field images revealed a characteristic ring shape on several of the Lamp1-EGFP positive structures (Supplementary Fig. 1a). The ring-shaped objects were usually between 1.2 and 0.9 µm enclosing a hollow space of varying size, consistent with the dimensions of lysosome organelles (Supplementary Fig. 1 b and c). Analyzing the intensity of moving objects usually introduce error on the imaging data and requires correction. As lysosomes are moving objects inside the living cell, we first evaluated the relative mobility of Lamp1-EGFP labeled membranes ^27^. By computing mobility maps we determine that at 20°C Lamp1-EGFP positive structures are relatively immobile during the two-minute window required for our recordings, eliminating the need for further correction of motion (Supplementary Fig. 1d).

An important contributor to the lysosomal membrane potential is the pH gradient (∆µH^+^), maintained by the vesicular proton pump (v-ATPase) ^28^. Thus, perturbation of the ∆µH_+_ would provide a simple approach to test our ability to measure Ψ_ly_ using hVoS_org_ in living cells. We first asked whether alkalinization of the lysosomal lumen can be induced directly by ammonium ^15,28,29^. As expected, ammonium incubation (10 mM) causes a rapid and strong change in luminal pH, observed as an increase of fluorescence signal measured from the pH sensor pHluorin, which we localized to the lysosomal lumen (Fig. 2 b and d). Next, we repeated the experiment using Lamp1-EGFP in the presence and absence of DPA (Fig. 2 c and d). It has been estimated that 20 mM ammonium in the extracellular solution will depolarize the lysosomal membrane in about 40mV ^15^. Accordingly, in the presence of DPA, ammonium quenches about 20% of GFP’s fluorescence indicating a voltage-dependent DPA transit within the lysosomal membrane (Fig. 2d). Such quenching is absent when the voltage sensor is not present. On the contrary, a modest increase in EGFP’s fluorescence can be detected after the ammonium treatment in the absence of DPA (Fig. 2d). This could be explained because NH4^+^ will alkalinize not only cellular compartments but also the cytoplasm. It has been reported that 20 mM ammonium in the external solution will cause a change in cytoplasmic pH of about 0.3 pH units ^30^. Given the high buffer capacity of the cellular cytoplasm, we reasoned that normal fluctuations in cytoplasmic pH would not contaminate our membrane potential measurements. Under our imaging conditions (i.e. GFP facing the cytosol), a negative deflection of the fluorescence signal is caused by the redistribution of DPA molecules to the less negative outer leaflet of the lysosomal membrane (Fig. 2d), and we interpret this as depolarization of the lysosomal membrane. For the case of single membrane organelles (i.e. ER, lysosomes, endosomes, and golgi) ΔΨ is calculated by Vm_org_= V_cytosol_-V_lumen_ ^31^, therefore our results indicate that alkalinization of the lysosome lumen causes a rapid depolarization of the organelle’s membrane as reported before ^15^.

To further examine whether hVoS_org_ is capable of following rapid changes in Ψ_ly_, we used the optogenetic tool Lyso-pHoenix. Lyso-pHoenix is a large protein sensor composed by mKate at the cytoplasmic N-terminal domain followed by the light activated proton pump Arch, and pHluorin on the luminal C-terminal domain of the protein ^26^. The sensor targets the lysosome via a Lamp3 destination signal (Supplementary Fig. 2a). The light-dependent activity of Arch is clearly visible in the lysosome upon v-ATPase inhibition by Bafilomycin A1 (Baf), which is used to initially prevent proton influx into the lysosomal lumen ^26^. After 20 minutes of incubation with Baf (300 nM) and DPA (4 µM), light pulses (10 mW s^2^ µm^2^; 560 nm) delivered at 0.3 Hz induces the activation of Arch, causing lysosomal acidification as monitored by fluorescence of pHluorin (Supplementary Fig. 2b). This acidification correlates with a simultanous increase in the fluorescence signal of the otherwise voltage-insensitive mKate protein, which indicates a redistribution of DPA towards the luminal leaflet of the lysosomal membrane, away from mKate (Supplementary Fig. 2b and c). The observed increase in mKate fluorescence suggests a repolarization of the lysosomal membrane. Taken together, our approach not only demonstrates the ability to follow the amplitude and kinetics of changes in Ψ_ly_ but also confirms the importance of luminal pH in setting the resting potential of lysosomes.

## Calibration of the hVoS_org_ signal

The transport of ions into the lumen of the lysosome depends on the electrochemical gradient. At rest, the electrochemical gradient of lysosomes can be roughly separated in two components – the chemical gradient of protons (H^+^) (∆pH_ly_), and the lysosomal membrane potential (Ψ_ly_) ^32^. Ammonium treatments provides us with a rough estimate of the voltage versus fluorescence response of our probe. To better control over the voltage across the lysosomal membrane, we performed an *In-cell* calibration of hVoS_org_ by using potassium (K^+^) as the only permeating ion. Gently digitonin-permeabilized cells were incubated with nigericin, an antiporter of H^+^ and K^+^ (to dissipate ∆pH_ly_), and the K^+^-selective ionophore valinomycin, to have control of Ψ_ly_ simply by changing the K^+^ concentration in the extracellular solution now in contact with the external membrane of the lysosome (Fig. 3 a and b). By doing this, we observed a linear of the response up to 120 mV (positive inside). Considering the signal-to-noise ratio we estimated the limit of detection to be 0.9±0.4 % of ΔF/F, corresponding to about 8mV (Fig. 3b).

**Figure 3.**
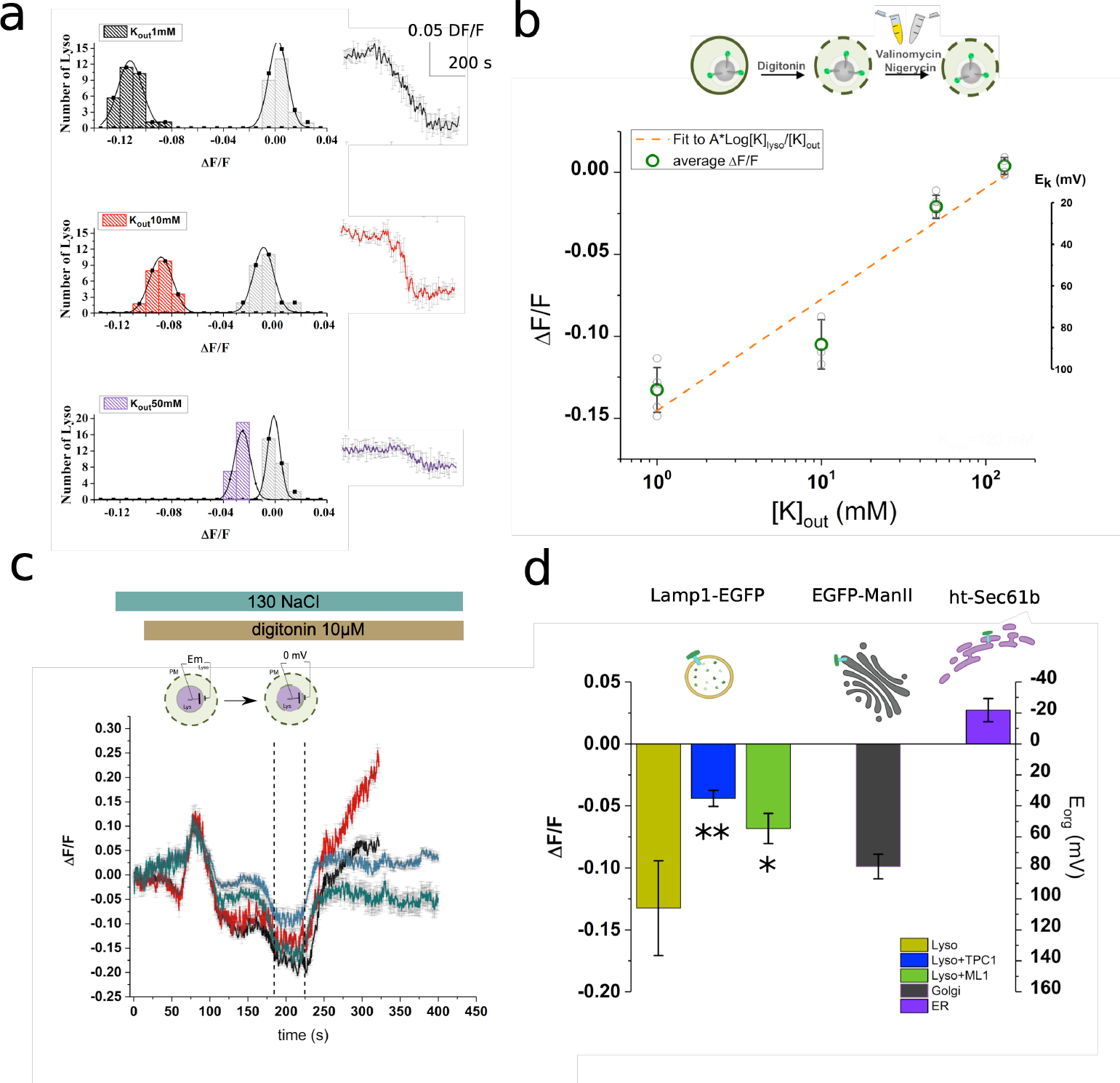
Organellar membrane potentials are measured at rest using hVoS_org_. **(a)** Representative traces and histograms used during the calibration procedure. Data represents one single experiment in which we observed the paired response of more than 20 lysosomes per condition. **(b)** *In-cell* calibration of hVoS_org_ by potassium clamp. The cartoon on top indicates the steps of the procedure, which included gentle permeabilization and incubation of ionophores. Averaged ΔF/F values (green circles) from 5 independent experiments (gray circles) were plotted versus the lysosomal external concentration of potassium. The experimental values were fitted to a Nernst equation as indicated on the top left corner, where A corresponds to (RT/zF); z= 1, F, R, and T have their usual meanings. The concentration of potassium when ΔF/F is zero was estimated to be 120 mM, indicating that the equilibration procedure was effective. **(c)** Representative traces used to estimate the resting potential. Cells were incubated in normal extracellular ringer and exposed to 10 µM digitonin as indicated on top. The cartoon indicates the expected values of membrane potential. During the procedure the cell experience two sequential steps of permeabilization (plasma membrane and lysosomal membrane), indicated in the cartoon. The second permeabilization step brings Ψ_ly_ to zero (indicated by dashed vertical lines) before the final disruption of the lysosomal membrane. **(d)** Resting membrane potential of Lysosomes, Golgi, and ER. Error bars indicate Mean ± SD; **P < 0.01 (one-way ANOVA with Bonferroni post hoc test).

By simply analyzing changes in ΔF/F of lysosomes transiting from resting to a condition where the Ψ_ly_ should be close to zero (i.e. digitonin permeabilization of the lysosomal membrane) we estimated that the resting potential of the lysosome is about 115 ±22 mV (lumen positive; n=7) (Fig. 3 c and d; supplementary movie 1). When observed in more detail, we observe that the Lamp1-positive structures on the periphery appear to have a smaller Ψ_ly_ at rest (Supplementary Fig. 3). We estimated that Ψ_ly_ ranges from 60 to 110 mV (positive inside) and would not be unreasonable to propose that these differences can be explained by the pH gradient observed in lysosomes during maturation ^33^.

We used the same experimental approach to examine the resting transmembrane potential of Golgi (Ψ_go_) and ER (Ψ_ER_) endomembranes. The Golgi marker manosidase II fused to EGFP (EGFP-ManII) reported 79±6 mV (positive inside; n=4) and the ER marker Sec61b fused to a HaloTag ^34^ (ht-Sec61b + Janelia Fluor 556) reported −25±9 mV (negative inside; n=3) (Figure 3d). Measuring the transmembrane potential of trans Golgi network has been elusive and previously estimated close to zero mV ^35^. By localizing hVoS_org_ to the Golgi membrane we present the first direct measurement of resting membrane potential of this compartment in living cells. Moreover, the confirmation that HaloTag can be used in combination with DPA opens many possibilities for multiplexing intracellular voltage signals and to explore more subtle aspects of lysosomal and cellular physiology ^22,23,36,37^.

### Modulation of lysosomal membrane potential by TRPML1 and TPC channels

To explore the contribution of known ion channels that are native in lysosomal membranes, we used hVoS_org_ to estimate Ψ_ly_ at rest with overexpression of TPC1 and TRPML1. In both cases a smaller quenching of the GFP signal was obtained, suggesting that the lysosomal membrane is less polarized upon overexpression (Fig. 3d). This observation suggests that the overexpressed channels are active and that their activity cannot be compensated efficiently by the v-ATPase or other lysosomal control mechanisms. TPC1 seems to be particularly effective on collapsing the resting potential of the lysosome. This suggests that the intrinsic voltage-sensitivity of the channel might create a positive-feedback loop when is not well compensated.

The mammalian target of rapamycin (mTOR) is a kinase that integrates intracellular level of nutrients, the energetic state, and growth factor signaling in higher eukaryotes ^38^. Starvation and/or rapamycin treatment (a general mTOR inhibitor) induce a robust electrical response of the endosome/lysosome (EL) vesicular compartment, linking mTOR signaling and TPC sodium channels ^39^. Moreover, the activity of mTORC1 has been associated to the activity of the SLC sodium-coupled amino acid transporter and also to the activity of the lysosomal v-ATPase ^40,41^. Consistent with the notion that the mTOR-signaling network is associated to lysosomal electrical activity ^42^, we observed that incubations with rapamycin (5 µM) evoked a strong and transient depolarization in intact lysosomes of living cells (Fig. 4; supplementary movie 2). The depolarization of the lysosomal membrane is in agreement with previous reports showing a Na^+^ efflux from the lumen of enlarged endolysosomes into the cytosol ^6,39^. It is worth noticing that even in the presence of rapamycin a late repolarization component is clearly visible, which may correspond to a voltage dependent component that could be fulfilled by BK channels present at the lysosomal membrane (Fig. 4c)^43^. Additionally, we attempt to multiplex the signals by co-expressing Lamp1-EGFP and ht-Sec61b.

**Figure 4.**
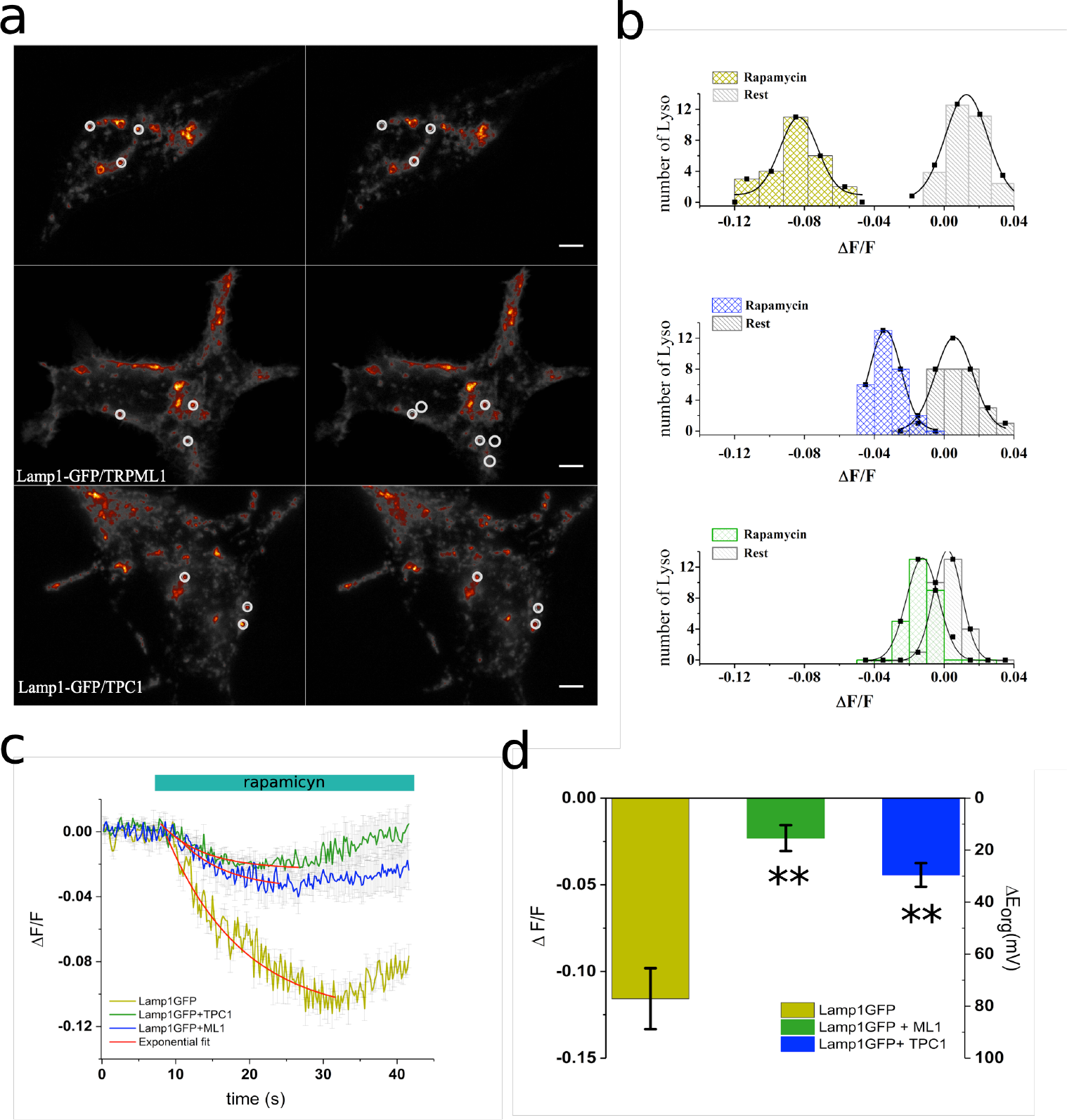
hVoS_org_ can detect a rapamycin-induced, lysosome-specific electrical response. **(a)** Pseudo color images of HEK293 cells transiently expressing Lamp1-EGFP alone (*top panels*) or co-expressing either TRPML1 or TRPC channels (*middle and bottom panels respectively*). Bar = 5 µm. **(b)** Representative traces and histograms used during the calibration procedure. Data represents one single experiment in which we observed the paired response of more than 25 lysosomes per condition. (c) Representative traces of rapamycin-induced depolarization of lysosomes in the different conditions. Red traces correspond to the fit to a first order exponential decay function. **(d)** Changes in the lysosomal membrane potential in the presence of the mTOR inhibitor rapamycin (5 µM). Error bars indicate Mean ± SD; **P < 0.01 (one-way ANOVA with Bonferroni post hoc test).

Simultaneous imaging of ΔΨ_ly_ and ΔΨ_ER_ was performed in response to rapamycin and followed by digitonin permeabilization, showing a clear temporal separation of responses and the ability to space-resolve the signals from both compartments (supplementary movie 3).

We then compared the averaged response of lysosomes to rapamycin in cells expressing Lamp1-EGFP alone or co-expressed with either hTPC1 or hTRPML1 ion channels (Fig. 4a). We calculated that the rapamycin-dependent depolarization dissipates the lysosomal resting potential by 85 ± 4 mV (n=4) (Fig. 4 c and d), which corresponds to nearly 75% reduction of Ψ_ly_ at rest. In contrast, the overexpression of both TPC1 and TRPML1 channels showed a lower effect on Ψ_ly_ (30 ± 6 and 15 ± 4 mV respectively; n=3) (Fig. 4 c and d). The time course of the response can be described by a single exponential decay with a characteristic time constant that is significantly different between normal lysosomes (12.3±3.3 s; n=4) and those with lysosomal channels overexpressed (p<0.01; Fig. 4c). The kinetics of the lysosomal response is shifted towards smaller values for both TPC1 (4.8±1.6 s; n=3) and TRPML1 transfected cells (5.7±1.2 s; n=3) (Fig. 4c). The acceleration observed in the response when TPC1 channels are overexpressed suggests that they are not close to the maximal open probability under overexpression conditions ^6^.

Although the smaller dissipation of the voltage gradient correlates well with the ability to set a more depolarized resting potential, the acceleration on depolarization kinetics together with the appearance of a late seemingly voltage-dependent component, suggests to us that we cannot work under the assumption that the lysosome membrane operates as a simple resistor-capacitor circuit. Thus, our results suggest that the overexpression of ion channels that are residents of the lysosome affect the resting potential and in doing so the dynamic response of the lysosome.

## Discussion

The membrane potential is a major regulator of electrogenic transport across membranes. Therefore, organelle function and by extension the metabolic state of the cell, is modulated by the membrane potential of organelles. Lysosome membranes contain several voltage-gated ion channels and transporters, suggesting that a fine-tuning of lysosomal function is modulated by the membrane potential ^22^. Imaging ΔΨ_ly_ have been successfully accomplished in the past by using a combination of potentiometric fluorescent dyes (i.e. oxonol derivatives) forming a FRET pair with fluorophores that are *preferentially* segregated to the lysosome membrane ^15^. However, the high density and variety of membrane-bound organelles make impossible to isolate the individual contribution of endosomes, lysosomes or phagosomes when a FRET pair is formed by hydrophobic dyes, imposing restrictions on spatial resolution and targeting. When a FRET system is set between a pair of light emitting probes, spectral properties have to be controlled to avoid optical leak and bleed-through. As the FRET pair used here consists of a fluorescent protein donor and a colorless acceptor, there is no need for bleed-through correction. By changing the paradigm, providing a colorless FRET acceptor reaching intracellular membranes combined with well-tested organelle markers, our single wavelength excitation method contributes on solving the problem of targeting the FRET pair to specific intracellular compartments together with providing a larger set of optical markers to be used. Moreover, DPA have the advantage of acting as FRET acceptor for fluorescent proteins in all visible range making an ideal sensor for optical multiplexing of organelle’s activity.

By targeting hVoS_org_ we effectively measure changes in membrane potential resolved in time and space within living cells (Fig. 5), an experimental feat that has not been previously possible. The present hVoS_org_ approach not only allows recording of membrane potentials in intact cells but demonstrate to be robust enough measure the elusive resting potential of intact individual sub-cellular structures that include lysosomes, Golgi, and ER.

**Figure 5.**
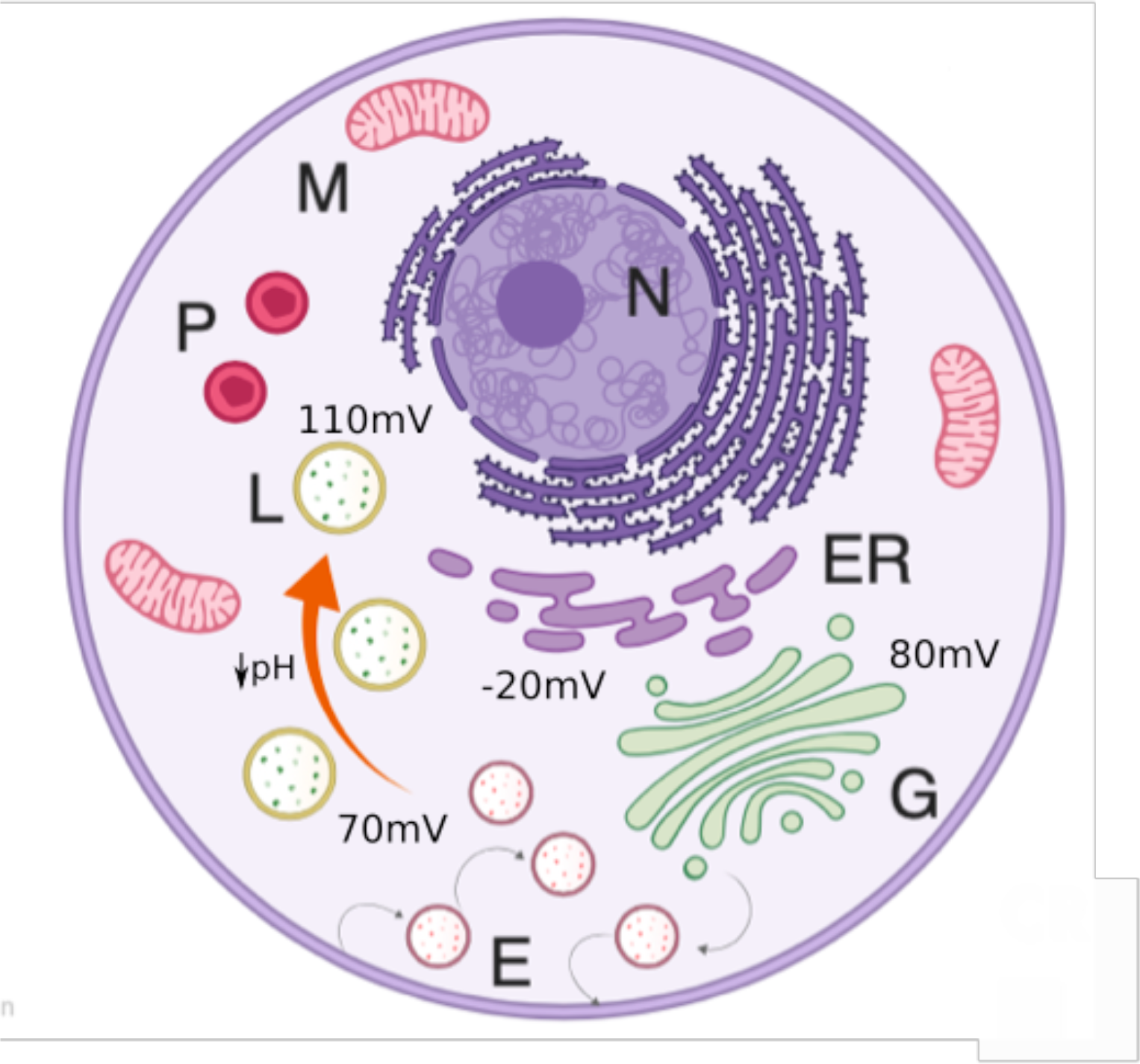
Schematic model of organellar membrane potential. Under normal conditions, lysosomes (L, yellow) and golgi (G, green) compartments have a relatively large and positive resting potential that contrast with a more modest and negative inside potential observed for ER (purple). Pumping of protons into the lysosomal lumen by the V-ATPase leads to acidification, causing a more hyperpolarized membrane potential in the mature lysosome (orange arrow).

We also showed the ability of performing simultaneous measurements from different intracellular membranes, demonstrating the capacity of hVoS_org_ to provide new insights for cell biologists into whether organelle-localized signals are modulated as a result of changes in organellar membrane potentials. We foresee that combinations of sensors having different spectral properties, targeted to distinct sub-cellular compartments, will allow for detailed space-time correlations of organelle’s activity in living cells^37,44^.

## METHODS

### Cell culture and clones

HEK293 cells were cultured in DMEM supplied with 10% FBS. Cells were plated in poly-l-lysine coated coverslips and transfected using lipofectamine 2000 (Invitrogen). Recordings were made 24-36 hours after transfection. Lamp1GFP was a gift from Patricia Burgos (Universidad Austral de Chile), hTPC1 was a gift from Dejian Ren (University of Pennsylvania), TRPML1 was a gift from Kirill Kiselyov (University of Pittsburg), halo-Sec61b and sapphire-manosidase II were a gift from <u>Jennifer Lippincott-Schwartz (Janelia</u> Research Center), farsenylated EGFP was a gift from Baron Chanda, hVoS (Addgene plasmid # 45282) was a gift from Meyer B. Jackson, Lyso-pHluorin (Addgene plasmid # 70113) and Lyso-pHoenix (Addgene plasmid # 70112) were a gift from Christian Rosenmund, and mTurquoise2-ER (Addgene plasmid # 36204) was a gift from Dorus Gadella. EGFP-ManII was obtained by PCR amplification of Sec61b from sapphire-ManII and introducing the amplified fragment in pEGFP-C1 vector.

### Reagents

Dipicrylamine sodium salt was obtained from Biotium Inc. (Fremont, CA). Rapamycin, DMSO, and ammonium chloride were obtained from Sigma-Aldrich. Valinomycin and nigericin were obtained from Tocris Bioscience (Bristol, UK). Standard salts used for solutions were obtained from Merck.

### Solutions and drug delivery

Dipicrylamine sodium salt (DPA) was prepared fresh at 20 mM stocks in DMSO and used at a final concentration of 4 µM. Extracellular solution contained NaCl (140 mM), KCl (5 mM), Ca Cl_2_ (2mM), Mg Cl_2_ (2mM), Glucose (5 mM), HEPES (8 mM) at pH 7.34. Cells were incubated with DPA for at least 10 minutes before start the experiments. In all the experiments the ringer solution contained DPA (4 µM) and DMSO (0.15 % v/v). At all times the concentration of both DPA and DMSO remained constant in the external solution. The drug-delivering pipette was placed close to the cell using a mechanical manipulator (Narishige, Tokio, Japan) and was pressure-ejected using a microliter syringe (Supplementary Figure 4). To make the voltage versus fluorescence calibration curve cells were first gently permeabilized with digitonin (10 µM, 3 min) in a solution containing KCl (130 mM), NaCl (10 mM), Ca Cl_2_ (2mM), Mg Cl_2_ (2mM), MgATP (5 mM), Glucose (5 mM), HEPES 8mM at pH 7. 4. Nigericyn and Alamethicyn were delivered to the extracellular solution in contact with the lysosomal membrane after permeabilization. Prior solution exchange, lysosomes were equilibrated with the 130 mM K^+^ buffer for 5 to 10 minutes, in the presence of antibiotics.

### Image acquisition. Voltage Imaging

An Orca Flash 4.0 CMOS camera (Hamamatsu Photonics, Japan) was used to image fluorescence. Acquisition was performed in streaming mode sampling at 20-10Hz without binning. For some experiments an electronic shutter (Uniblitz, VA Inc., Rochester, NY) was used. The camera was mounted on an Olympus IX71 microscope. Images were taken under normal epifluorescence, using a water immersion objective (60x, N.A.=1.3). A 473 nm (Melles Griot, Carlsbad, CA) and 532 nm (LaserGlow, Toronto, Canada) diode pump lasers were transmitted via the rear illumination port of the microscope and reflected to the sample by a double dichroic mirror with reflection bands at 473-490 nm and 530-534 nm and transmission bands at 500-518 nm and 550-613 nm (Semrock, Rochester, NY) (Supplementary Figure 4). *Space correlated imaging*. The optical system described above was used in combination to a DualView system (Photometrics, Tucson, AZ), splitting the emission signal in two channels (490/40 and 600/70 nm) that are focused simultaneously on the CMOS chip (Supplementary Figure 4). All image acquisition was controlled by micro-manager (Open Imaging, San Francisco, CA).

### Signal analysis. Colocalization

After background subtraction, 20 frames of space-correlated images were averaged and overlay images were produced further over imposed on Cairn optosplit plugin for ImageJ. To evaluate pixel colocalization we calculate Pearson’s coefficient using JACoP plugin for ImageJ ^45^. *Mobility maps*. We performed the calculation following the protocol in Brauchi et al. 2008. The fluorescence signal was lower-threshold over 2 standard deviations above the mean of the camera noise. The upper threshold was set identify the desired lysosomal-shaped objects. Threshold images were converted into binary format events. The mobility function was calculated for each pixel of the image sequence. *Fluctuation of fluorescence*. To calculate fluorescence kinetics on pHluorin and DPA-GFP experiments, regions of interest (ROIs) were established on doughnut shaped, Lamp1-positive spots having a size between 50 and 200 nm (5-20 pixels), located on regions of low mobility. Several regions of interest (ROIs) per cell were selected from background-subtracted stacks. All together shape, size, mobility, and localization helped us to define the ROI set that was measured on each cell. After selection, fluorescence time course was recorded for each ROI. Numerical data was treated and plotted using OriginPro (OriginLab corporation, Wellesley Hills, MA). Fractional fluorescence change was calculated according to: ΔF/F = ((F_n –_ F_0_) / F_0_), where, F_n_ is the corrected fluorescence at frame n, and F_0_ corresponds to the average of at least 10 frames of the base line fluorescence. Baseline fluorescence corresponds to the steady state GFP signal after DPA equilibration. Once ΔF/F was calculated bleaching was corrected by using a single exponential function fitting the decay of fluorescence at baseline. Data was often filtered by a FTT filter (implemented in OriginPro) with a cut off 1/4^th^ of the sampling frequency.

### Statistics and figure preparation

Voltage measurements. Individual cells on one day of transfection contribute with several lysosomes (in the order of 10 to 25 per cell), which are considered as replicates. Therefore the histograms presented corresponds to n = 1 and contains information from several individual lysosomes. The final statistical procedure was done by performing one-way ANOVA comparing the differences observed in several independent experiments per condition. A p-value of 0.01 was considered significant and a Bonferroni post-test was performed for all pairwise comparisons tests. Plotting the mean value of the signal generated histograms that were used to present data. The histograms were fitted to a Gaussian distribution function to evaluate normality (also evaluated by Kolmogorov-Smirnov test implemented in Origin). The statistical analysis were computed in Microcal OriginPro ver9 (OriginLab) Figures were prepared using Microcal OriginPro ver9 and ImageJ.

## Acknowledgments

We thank David E Clapham (HHMI) for the use of his microscopes and support; Kirill Kiselyov (U Pittsburg) for his helpful insights, and Alexandria Miller (HHMI) for her critical reading of the manuscript. This work was supported by Anillo Cientifico #ACT1401. SB is part of UACh Program for Cell Biology. MiNICAD is a Millennium Nucleus supported by Iniciativa Científica Milenio, Ministry of Economy, Development and Tourism, Chile

