## Supplementary figures and legends for "Imaging the Electrical Activity Of Organelles in living cells"

SUPLEMENTARY MATERIAL

FIGURES and LEGENDS.

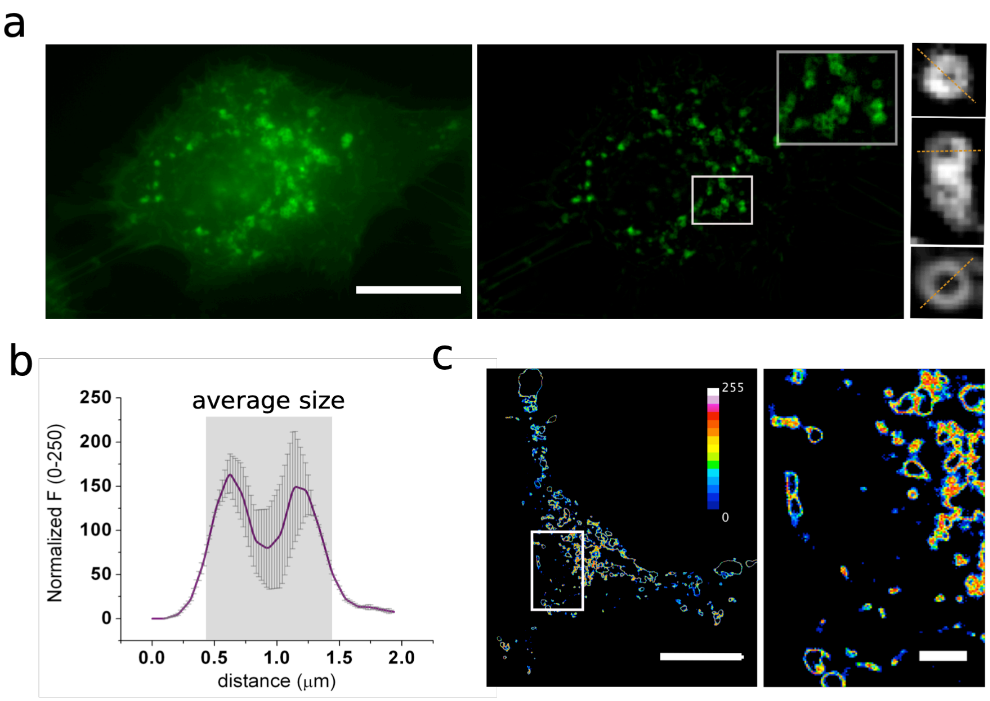


SFigure 1. Matamala et al.

**SF1**. Representative images of the sub cellular structures identified as lysosomes. **(a)** Raw image of a transiently transfected HEK293 cell expressing Lamp1-EGFP (*left*). FFT filtered and background subtracted image (*middle*). Individual lysosomes can be easily identified. Inset corresponds to a 2x2 μm zoom of the highlighted area. Average of 10 frames allow obtaining better resolved single lysosome structures (*right*). The orange dotted line indicates the axis used to extract the intensity of the Lamp1 positive objects. **(b)** Intensity versus distance plot showing the average size of lysosomes in the study. Grey values were collected along orange lines as indicated in a. The average size of the doughnut shaped objects was 112 ± 33 nm (n=26). error bars correspond to SEM**. (c)** Mobility map of a cell expressing Lamp-EGFP (*left*; Scale bar, 10 μm). The white square indicates the region zoomed on the right panel (Scale bar, 2 μm). Red shades indicate zones with high motion and black represent regions containing immobile particles.

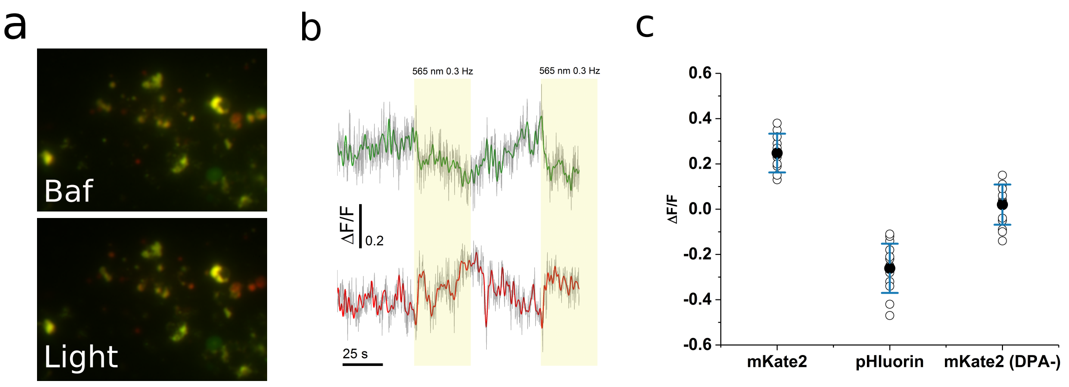


SFigure 2. Matamala et al.

**SF2**. Acidification of the lysosome hyperpolarize lysosomal membrane. **(a)** Images of HEK-293 cells expressing LysopHoenix after the treatment with bafilomycin A1 (*top*) and during light stimulation (*bottom*). **(b)** Simultaneous traces of pH (green) and membrane voltage (red) acquired during light stimulation (indicated in yellow). **(c)** Average values of ΔF/F for the change in membrane voltage (mKate2), luminal pH (pHluorin) and the control condition in the absence of DPA.

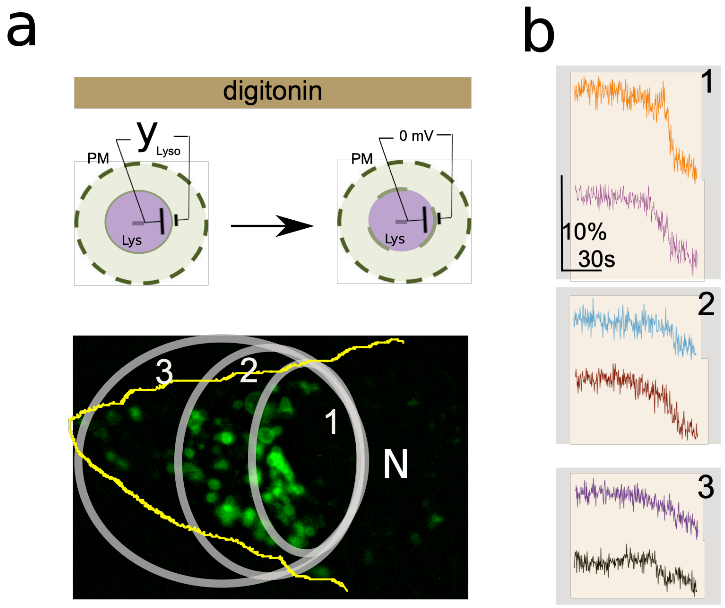


SFigure 3. Matamala et al.

**SF3.** Peripheral lysosomes are less depolarized. **(a)** Image of a cell transiently expressing Lamp1-EGFP. The yellow line indicates the cell membrane and N denotes nucleus. Concentric semi circular areas were defined and individual lysosomes on these areas were measures and compared. The cartoon on top depicts the experimental procedure. Digitonin permeabilization of the lysosomal membrane will dissipate all the memebrane potential at each individual structure. **(b)** Representative traces obtained from single lysosomes. The traces are grouped according to the areas defined in a.

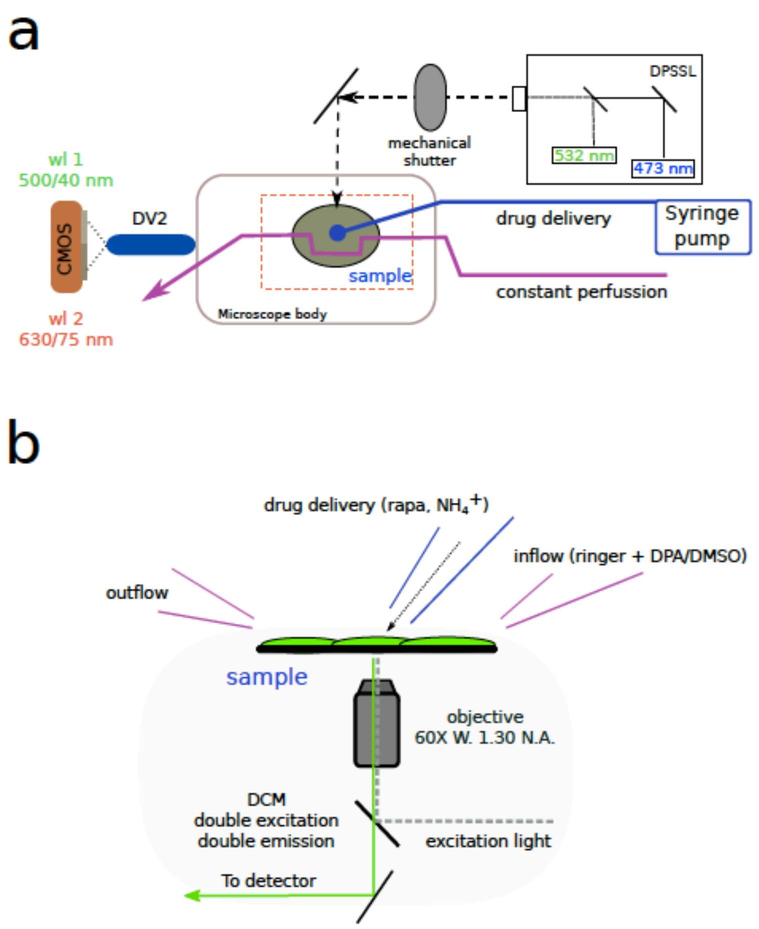


**SF4.** Optical layout and perfusion. **(a).** Schematic representation of the imaging and perfusion setup used in this work. The constant perfusion (purple line) and drug delivery system (blue line) are critical to maintain reproducibility. **(b).** The ringer solution being perfused contained DPA and DMSO to reduce variability due to fluctuations on vehicle concentration or DPA. Chemicals were applied directly to the cells via a syringe pump.

**Movie1.** $\Delta$F/F signal in pseudo color for a cell expressing $hVoS_{org}$ and treated with digitonin. Digitonin was added in frame 10.

**Movie2.** $\Delta$F/F signal in pseudo color for a cell expressing $hVoS_{org}$ and exposed to rapamycin. Rapamycin was added in frame 10.

**Movie3.** $\Delta$F/F signal in pseudo color for a cell expressing double $hVoS_{org}$ (i.e. Lamp1-EGFP and ht-Sec61b) exposed to rapamycin and digitonin. Rapamycin addition is indicated by a red dot on the upper right corner. Digitionin addition is indicated by a white dot on the upper right corner. While lysosomes darken while depolarizing, ER gets brighter upon depolarization.
